# SPINT2 inhibits proteases involved in activation of both influenza viruses and metapneumoviruses

**DOI:** 10.1101/752592

**Authors:** Marco R. Straus, Jonathan T. Kinder, Michal Segall, Rebecca Ellis Dutch, Gary R. Whittaker

**Author notes:** Address correspondence to Rebecca E. Dutch, or Gary R. Whittaker. M.R.S and J.T.K contributed equally to this work.

## Abstract

Viruses possessing class I fusion proteins require proteolytic activation by host cell proteases to mediate fusion with the host cell membrane. The mammalian SPINT2 gene encodes a protease inhibitor that targets trypsin-like serine proteases. Here we show the protease inhibitor, SPINT2, restricts cleavage-activation efficiently for a range of influenza viruses and for human metapneumovirus (HMPV). SPINT2 treatment resulted in the cleavage and fusion inhibition of full-length influenza A/CA/04/09 (H1N1) HA, A/Aichi/68 (H3N2) HA, A/Shanghai/2/2013 (H7N9) HA and HMPV F when activated by trypsin, recombinant matriptase or KLK5. We also demonstrate that SPINT2 was able to reduce viral growth of influenza A/CA/04/09 H1N1 and A/X31 H3N2 in cell culture by inhibiting matriptase or TMPRSS2. Moreover, inhibition efficacy did not differ whether SPINT2 was added at the time of infection or 24 hours post-infection. Our data suggest that the SPINT2 inhibitor has a strong potential to serve as a novel broad-spectrum antiviral.

## Introduction

Influenza-like illnesses (ILIs) represent a significant burden on public health and can be caused by a range of respiratory viruses in addition to influenza virus itself (1). An ongoing goal of anti-viral drug discovery is to develop broadly-acting therapeutics that can be used in the absence of definitive diagnosis, such as in the case of ILIs. For such strategies to succeed, drug targets that are shared across virus families need to be identified.

One common cause of influenza, which shaped the term ILI, are influenza A viruses, including H1N1 and H3N2, that cross species barriers from their natural avian hosts and infect humans (2, 3). Novel emerging viruses such as H7N9 that has caused hundreds of deaths since its appearance in China in 2013 pose an additional concern (4). The World Health Organization (WHO) estimates that each year about 1 billion suffer from flu infections, 3 to 5 million people worldwide are hospitalized with severe illness and approximated 290,000 to 650,000 people die from the disease (5). Mortality rates can dramatically rise during influenza pandemics as observed with the Spanish flu of 1918, the Asian pandemic of 1957 and the Hong Kong pandemic of 1968 (3, 6). Vaccinations can provide an effective protection against seasonal and pandemic outbreaks but provide limited or no protection when viruses evolve and/or acquire mutations resulting in antigenically distinct viruses. This antigenic drift or shift requires that new vaccines be produced quickly and in vast amounts which can be problematic especially during pandemics. In addition to vaccines, several antiviral therapies have been applied to treat influenza A infections such as adamantanes acting as M2 ion channel blockers (amantadine, rimantadine) and neuraminidase inhibitors which block the cleavage of sialic acid in newly formed virions (oseltamivir, zanamivir) (7). However, several influenza subtypes including the most common H1N1 and H3N2 have emerged globally that are resistant against adamantanes and similar observations were made with respect to oseltamivir (7). More recently, baloxavir marboxil (BXM) which is an inhibitor of the influenza polymerase acidic subunit (PA) was approved as an antiviral therapy (8). Although it is fully effective against currently circulating influenza A and B viruses clinical trial studies have already shown that treatment with BXM selects for influenza virus with specific amino acid substitutions in PA resulting in reduced susceptibility to the drug (9, 10).

Another common cause of ILI are pneumo- and paramyxoviruses, including human metapneumovirus (HMPV), respiratory syncytial virus, and parainfluenza viruses. Clinical presentation of these viruses resembles many of the symptoms of influenza, where they cause significant morbidity and mortality as well as a large economic burden (11, 12). HMPV is ubiquitous, with nearly everyone infected by the age of 5 and reinfection is common throughout life, impacting children, the elderly and immunocompromised individuals (13–15). While HMPV is a common cause of ILI, there are currently no approved vaccines or antiviral therapeutics. Further research is needed to establish targets for intervention, and factors required for infection need to be examined in more detail.

Certain influenza viruses and HMPV appear to share common activating proteases. The influenza fusion protein hemagglutinin (HA) is synthesized as a precursor that needs to be cleaved by host cell proteases to exert its fusogenic activity (16, 17). Cleavage separates the precursor HA_0_ into HA_1_ and HA_2_, which remain associated via disulfide bonds and leads to exposure of the fusion peptide at the N-terminus of HA_2_ (16, 17). Low pathogenicity avian influenza (LPAI) usually possess a monobasic cleavage site which consists of 1 – 2 non-consecutive basic amino acids and which is generally cleaved by trypsin-like serine proteases such trypsin, as well as members of the type II transmembrane serine protease (TTSP) family including TMPRSS2, TMPRSS4, HAT (TMPRSS11D) and matriptase (16–18). In addition, some other proteases such as KLK5 and KLK 12 have been implicated in influenza pathogenicity (19). In humans these proteases are localized in the respiratory tract and therefore influenza infections are usually confined to this tissue. In contrast, highly pathogenic avian influenza (HPAI) viruses are defined by a polybasic cleavage site which consists of 6 – 7 basic residues allowing them to be activated by members of the proprotein convertase (PC) family such as furin and PC6 (20). These proteases are not confined to a specific tissue and dramatically increase the risk of a systemic infection.

Similar to influenza HA, HMPV requires the fusion protein (F) to be proteolytically processed at a single basic residue to generate the active, metastable form. Without this cleavage process, the F protein is unable to mediate viral entry into the target cell. However, to date, only trypsin and TMPRSS2 have been shown to effectively cleave HMPV F and other proteases have yet to be identified (21, 22).

In addition, other respiratory viruses have been reported to utilize similar proteases for activation, including SARS-CoV and MERS-CoV, demonstrating that targeting these proteases would inhibit multiple respiratory pathogens (23). The fact that proteolytic activation is such a crucial step for several respiratory viruses that predominately require a specific class of proteases makes these proteases a viable target for the development of novel antiviral therapies (24). Earlier studies described the administration of the serine protease inhibitor aprotinin to inhibit influenza replication and demonstrated that aprotinin successfully inhibited IAV activation and replication (25). However, when targeting host specific factors there are potential off target effects, and therefore the potential side effects of targeting host proteases requires further investigation. Hamilton et al. reported that the hepatocyte growth activator inhibitor 2 (HAI-2) effectively inhibited trypsin-mediated cleavage of H1N1 and H3N2 *in vitro* and *in vivo* (26). HAI-2 is encoded by the SPINT2 gene and hereafter we will also refer to the protein as SPINT2. SPINT2 is 225 KDa plasma membrane-localized serine protease inhibitor found in epithelial cells of various tissues including the respiratory tract and all major organs (27). In most tissues, SPINT2 co-localizes with matriptase suggesting a regulatory role of SPINT2 on matriptase-mediated cleavage events. However, the finding that SPINT2 is also expressed in brain and lymph node cells indicates that it might regulate other proteases than matriptase (27). Recent reports associated the physiological role of SPINT2 with the inhibition of human serine-type proteases such as matriptase, plasmin, kallikreins (KLK) and coagulation factor XIa (28–31). SPINT2 possesses one transmembrane domain and two kunitz-type inhibitor domains that are exposed to the extracellular space and which are believed to facilitate a potent inhibition of target proteases. Wu et al. recently described that the kunitz-type domain 1 of SPINT2 is responsible for matriptase inhibition (28). A major function of SPINT2 is its role as a tumor suppressor because down-regulation diminishes the prospect of survival of several cancers such as hepatocellular carcinoma, gastric cancer, prostate cancer or melanoma (32–35). However, SPINT2 was also associated with placenta development and epithelial homeostasis (36, 37).

A previous study from our lab described the effective inhibition of trypsin by SPINT2 resulting in dramatically reduced cleavage of influenza A HA using a model protease and subsequently reduced viral growth in cell culture and mouse studies (26). Here, we report that purified SPINT2 protein inhibits several host proteases found in the human respiratory tract, such as matriptase and TMPRSS2, that are relevant for the activation of influenza viruses currently circulating and causing significant disease outbreaks. To demonstrate broad applicability, we also tested the potential of SPINT2 to inhibit the activation of the fusion protein (F) from human metapneumovirus (HMPV), a member of the pneumovirus family. We confirm the original findings that HMPV F is proteolytically processed by trypsin and TMPRSS2. In addition, we found that HAT, KLK5 and matriptase were able to cleave F, but KLK12 could not. Our results show that SPINT2 can inhibit the activation of proteases that are responsible for the activation of influenza H1N1, H3N2 and H7N9 HA as well as HMPV F. In a cell culture model, we demonstrate that viral loads are significantly reduced in the presence of SPINT2 when infections were conducted with A/CA/04/09 and A/X31. Moreover, the application of SPINT2 24 hours post infection inhibited the activation of influenza A viruses with the same efficacy as when SPINT2 was added to cell culture medium at the time of infection. Thus, SPINT2 exhibits the potential to serve as a novel and efficient antiviral therapeutic to relieve patients from influenza A, human metapneumovirus, SARS-CoV and potentially other respiratory viruses that require these host factors for entry.

## Results

### SPINT2 inhibits recombinant human respiratory tract proteases that cleave HMPV F and HA cleavage site peptide mimics

Using a fluorogenic peptide cleavage assay that utilizes fluorogenic peptides mimicking the HA cleavage site we previously tested the ability of SPINT2 to inhibit proteases shown to cleave HAs from seasonal and pandemic influenza A strains that infected humans (38, 39). We found that certain HA subtypes such as H1, H2 and H3 are cleaved by a wide variety of human respiratory proteases while others such H5, H7 and H9 displayed more variability in cleavage by proteases and seemed less well adapted to proteases present in the human respiratory tract (39). Here, we extended our previous study and tested a peptide mimicking the cleavage site of the pneumovirus fusion protein of HMPV F using a variety of proteases known for their ability to cleave the peptide mimic (Table 1) (22, 40). When we tested the cleavage of a peptide mimicking the HMPV F cleavage site using trypsin, matriptase, KLK5, KLK12, HAT or plasmin we found that all proteases except KLK12 were able to proteolytically cleave the peptide (Table 1). However, the Vmax values for matriptase (9.24 RFU/min), KLK5 (5.8 RFU/min) and HAT (2.99 RFU/min) were very low compared to trypsin (135.2 RFU/min) suggesting that the three proteases have a low affinity interaction and processivity for HMPV F.

**Table 1:** Cleavage profile of the HMPV F A: fluorogenic peptide mimicking the cleavage site of HMPV F was incubated with the indicated proteases and cleavage was monitored by the increase of fluorescence at 390 nm. RFU = relative fluorescence units. StDev = Standard Deviation.

|  | Vmax<br>(RFU/min) | StDev |
| --- | --- | --- |
| Trypsin | 135.25 | 17.55 |
| Matriptase | 9.24 | 0.39 |
| KLK5 | 5.80 | 0.12 |
| HAT | 2.99 | 0.41 |
| KLK12 | 0.00 | 0.00 |
| Plasmin | 37.86 | 2.07 |

Next, trypsin, matriptase and KLK5 were selected for the SPINT2 inhibition assays as described below. For the SPINT2 inhibition assays, trypsin (which typically resides in the intestinal tract and expresses a very broad activity towards different HA subtypes and HMPV F) served as a control (41). In addition, furin was used as a negative control that is not inhibited by SPINT2. As none of the peptides used in combination with the aforementioned proteases has a furin cleavage site we tested furin-mediated cleavage on a peptide with a H5N1 HPAI cleavage motif in the presence of 500 nM SPINT2 (Supplementary Figure 1). We continued by measuring the Vmax values for each protease/peptide combination in the presence of different SPINT2 concentrations and plotted the obtained Vmax values against the SPINT2 concentrations on a logarithmic scale (Supplementary Figure 2). Using Prism7 software, we then determined the IC_50_ that reflects at which concentration the V_max_ of the respective reaction is inhibited by half. SPINT2 cleavage inhibition of a representative H1N1 cleavage site by trypsin results in an IC50 value of 70.6 nM (Table 2A) while the inhibition efficacy of SPINT2 towards matriptase, HAT, KLK5 and KLK12 ranged from 11 nM to 25 nM (Table 2A). However, inhibition was much less efficient for plasmin compared with trypsin (122 nM). We observed a similar trend when testing peptides mimicking the H3N2 and H7N9 HA cleavage sites using trypsin, HAT, KLK5, plasmin and trypsin, matriptase, plasmin, respectively (Table 2A). With the exception of plasmin, we found that human respiratory tract proteases are inhibited with a higher efficacy compared to trypsin. We expanded our analysis to peptides mimicking HA cleavage sites of H2N2, H5N1 (LPAI and HPAI), H6N1 and H9N2 that all reflected the results described above (Table 2A). Only cleavage inhibition of H6N1 HA by KLK5 did not significantly differ from the observation made with trypsin (Table 2A).

**Table 2:**
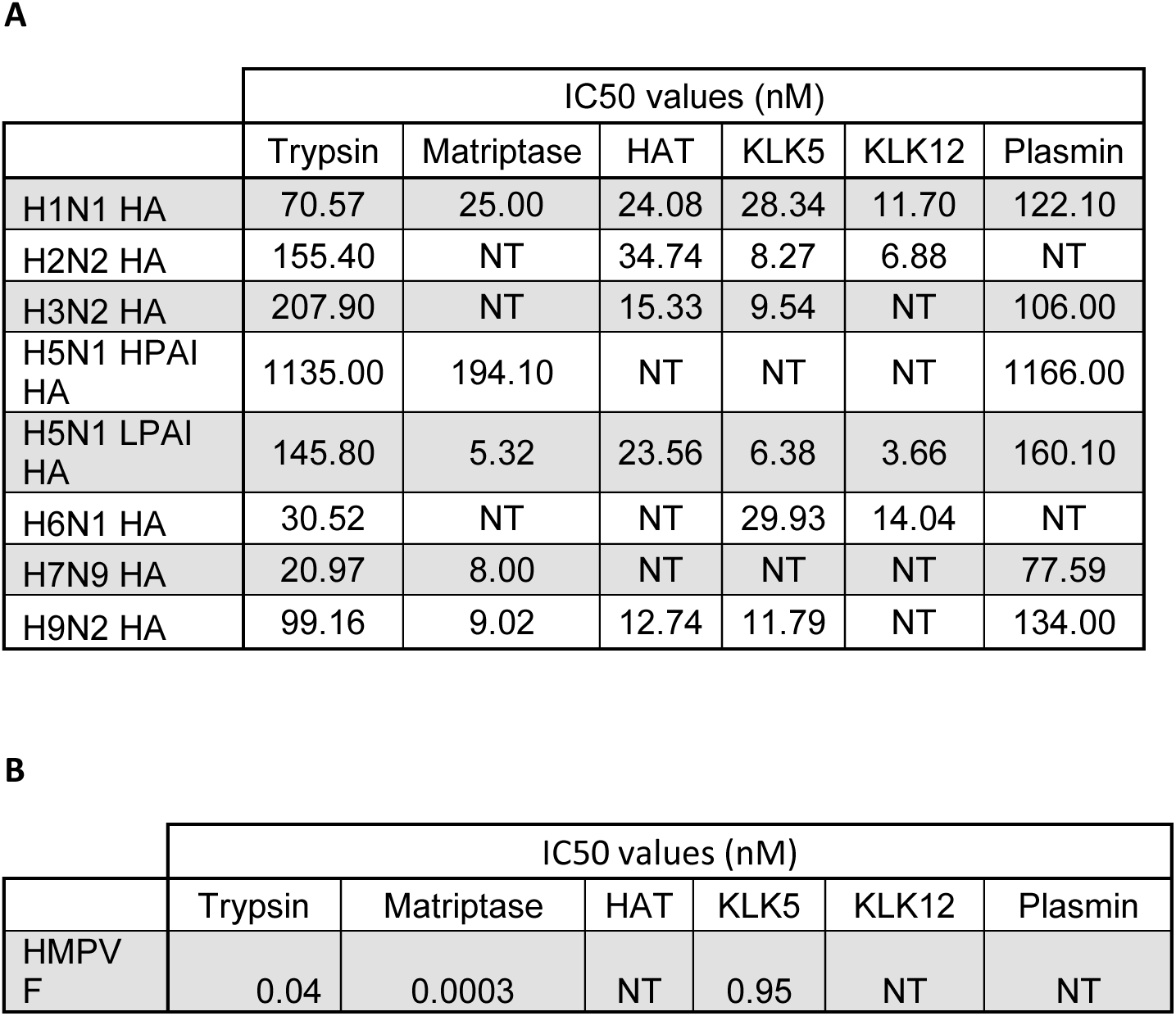
IC50 values of all protease/peptide combinations. Fluorogenic peptides mimicking the cleavage sites of A/CA/04/09 H1N1, A/Japan/305/1957 H2N2 HA, A/Aichi/2/68 H3N2 HA, A/Vietnam/1203/2004 H5N1 LPAI HA, A/Vietnam/1204/2004 H5N1 HPAI HA, A/Taiwan/2/2013 H6N1 HA, A/Shanghai/2/2013 H7N9 HA, A/Hong Kong/2108/2003 H9N2 HA and HMPV F were incubated with the indicated proteases and different SPINT2 concentrations. Cleavage was monitored by the increase of fluorescence at 390 nm and the resulting Vmax values were used to calculate the IC50 values as described in the “Material and Methods” section. (A) IC50 values of influenza A fluorogenic cleavage site peptide mimics. Concentrations are in nanomolar. (B) IC50 values of the HMPV F cleavage site peptide mimic. Concentrations are in picomolar. NT = Not tested.

When we tested the inhibition of HMPV cleavage by trypsin, matriptase and KLK5, SPINT2 demonstrated high inhibition efficacy for all three tested proteases with measured IC50s for trypsin, matriptase and KLK5 of 0.04 nM, 0.0003 nM and 0.95 nM, respectively (Table 2B).Compared to the IC50 values observed with the peptides mimicking influenza HA cleavage site motifs the IC50 values for the HMPV F peptide were very low.

### Cleavage of distinct full-length HA subtypes and HMPV F is efficiently inhibited by SPINT

Cleavage of peptides mimicking cleavage sites of viral fusion proteins do not always reflect the *in vivo* situation and requires-validation by expressing the full-length fusion proteins in a cell culture model to test cleavage and cleavage inhibition of the respective protease (39). However, before conducting these experiments we wanted to ensure that SPINT2 does not have a cytotoxic effect on cells. Therefore, 293T cells were incubated with various concentrations of SPINT2 over a time period of 24 hours. PBS and 500µM H_2_O_2_ served as cytotoxic negative and positive controls respectively. We observed a slight reduction of about 10-15 % in cell viability when SPINT2 was added to the cells (Figure 1).

**Figure 1:**
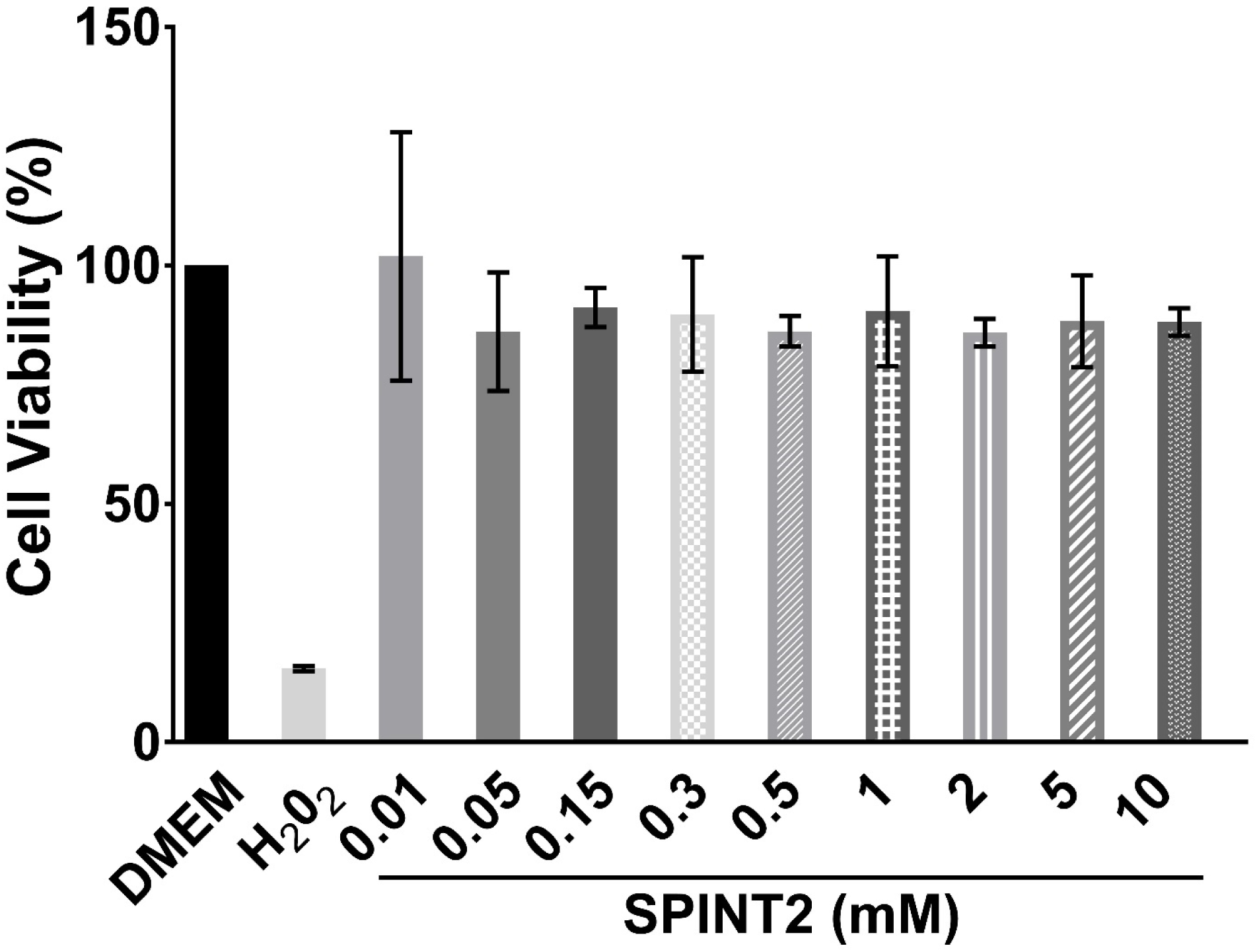
Cytotoxicity assay to evaluate the cytotoxic effect of SPINT2. 293T cells were incubated with indicated SPINT2 concentrations for 24 hours. DMEM and 500 µM H_2_O_2_ served as controls. After 24 hours cell viability was determined via a spectrophotometric assay.

To test SPINT2-mediated cleavage inhibition of full-length HA we expressed the HAs of A/CA/04/09 (H1N1), A/x31 (H3N2) and A/Shanghai/2/2013 (H7N9) in 293T cells and added recombinant matriptase or KLK5 protease that were pre-incubated with 10nM or 500nM SPINT2. Trypsin and the respective protease without SPINT2 incubation were used as controls. Cleavage of HA_0_ was analyzed via Western Blot and the signal intensities of the HA_1_ bands were quantified using the control sample without SPINT2 incubation as a reference point to illustrate the relative cleavage of HA with and without inhibitor (Figure 2A and B-D). Trypsin cleaved all tested HA proteins with very high efficiency that was not observed with matriptase or KLK5 (Figure 2B-D). However, H1N1 HA was cleaved by matriptase and KLK5 to a similar extent without and with 10nM SPINT2. 500nM SPINT2 led to a relative cleavage reduction of about 70% and 50% for matriptase and KLK5, respectively (Figure 2A and B). KLK5-mediated cleavage of H3N2 HA was reduced by about 10% when KLK5 was pre-incubated with 10nM SPINT2 and by about 60% when 500nM SPINT2 was used (Figure 2A and C). When we tested the cleavage inhibition of matriptase with H7N9 HA as a substrate we found that 10nM and 500nM SPINT reduced the cleavage to 40% and 10% cleavage, respectively, compared to the control. (Figure 2A and D). In contrast, 10nM SPINT2 had no effect on KLK5-mediated cleavage of H7N9 HA while 500nM reduced relative cleavage by approximately 70% (Figure 2A and D).

**Figure 2:**
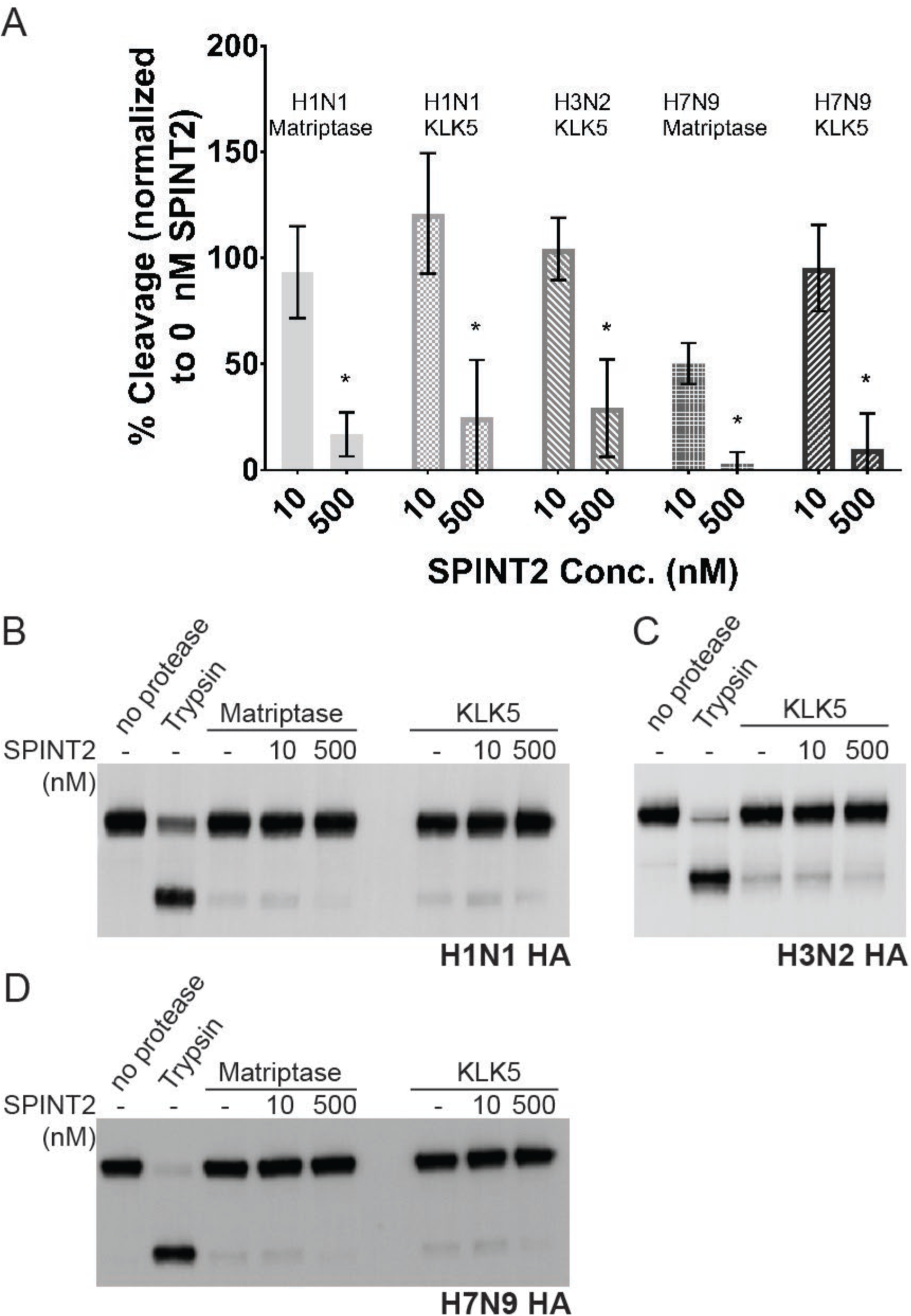
SPINT2 inhibits cleavage of HA protein expressed in 293T cells. Cells were transfected with plasmids encoding for the indicated HA and allowed to express the protein for ∼18 hours. The recombinant proteases were incubated for 15 minutes with the indicated SPINT2 concentrations and subsequently added to the cells for 10 minutes (trypsin) or 90 minutes (matriptase and KLK5). Western blots were performed and the HA_1_ band was quantified using ImageJ. (A) Quantification of the HA1 band comparing the signal intensity of the 0 nM SPINT2 samples against 10 nM and 500 nM SPINT2 of the respective HA/protease combination. Three independent experiments were carried out and the western blots of each experiment were analyzed. Quantifications were conducted as described in the methods section. (B – D) Western blots showing the cleavage of (B) A/CA/04/09 H1N1 HA by matriptase and KLK5 at different SPINT2 concentration, (C) A/Aichi/2/68 H3N2 HA by KLK5 at different SPINT2 concentration and (D) A/Shanghai/2/2013 H7N9 HA by matriptase and KLK5 at different SPINT2 concentrations. Statistical analysis was performed using a non-paired student’s t-test comparing the samples tested with 10 nM SPINT2 against the respective sample incubated with 500 nM SPINT2. Error bars indicate standard deviation. * indicates p = < 0.05.

In order to determine whether SPINT2 also prevented cleavage of HMPV F we first examined which proteases, in addition to trypsin and TMPRSS2, were able to cleave HMPV F. First, we co-transfected the full length TMPRSS2, HAT and matriptase with HMPV F in VERO cells. The F protein was then radioactively labeled with ^35^S methionine and cleavage was examined by quantifying the F_0_ full length protein and the F_1_ cleavage product. We found that TMPRSS2 and HAT were able to efficiently cleave HMPV F while matriptase decreased the expression of F, though it is not clear if this was due to general degradation of protein or lower initial expression. However, matriptase demonstrated potential low-level cleavage when co-transfected (Figure 3A and B). We then examined cleavage by the exogenous proteases KLK5, KLK12 and matriptase. Compared with the trypsin control, KLK5 and matriptase were able to cleave HMPV F, while KLK12 was not (Figure 3C and D). In agreement with the peptide assay, cleavage of HMPV F by KLK5 and matriptase was less efficient than for trypsin and both peptide, and full-length protein assays demonstrate that KLK12 does not cleave HMPV F. This also serves as confirmation that matriptase likely cleaves HMPV F, but co-expression with matriptase may alter protein synthesis, stability or turnover if co-expressed during synthesis and transport to the cell surface. Next, we tested SPINT2 inhibition of exogenous proteases trypsin, KLK5 and matriptase. We pre-incubated SPINT2 with each protease, added it to VERO cells expressing HMPV F and analyzed cleavage product formation. SPINT2 pre-incubation minimally affected cleavage at a concentration of 10nM but addition of 500nM SPINT2 resulted in inhibition of trypsin, KLK5 and matriptase-mediated cleavage of HMPV, similar to our findings for HA (Figure 3E and F).

**Figure 3:**
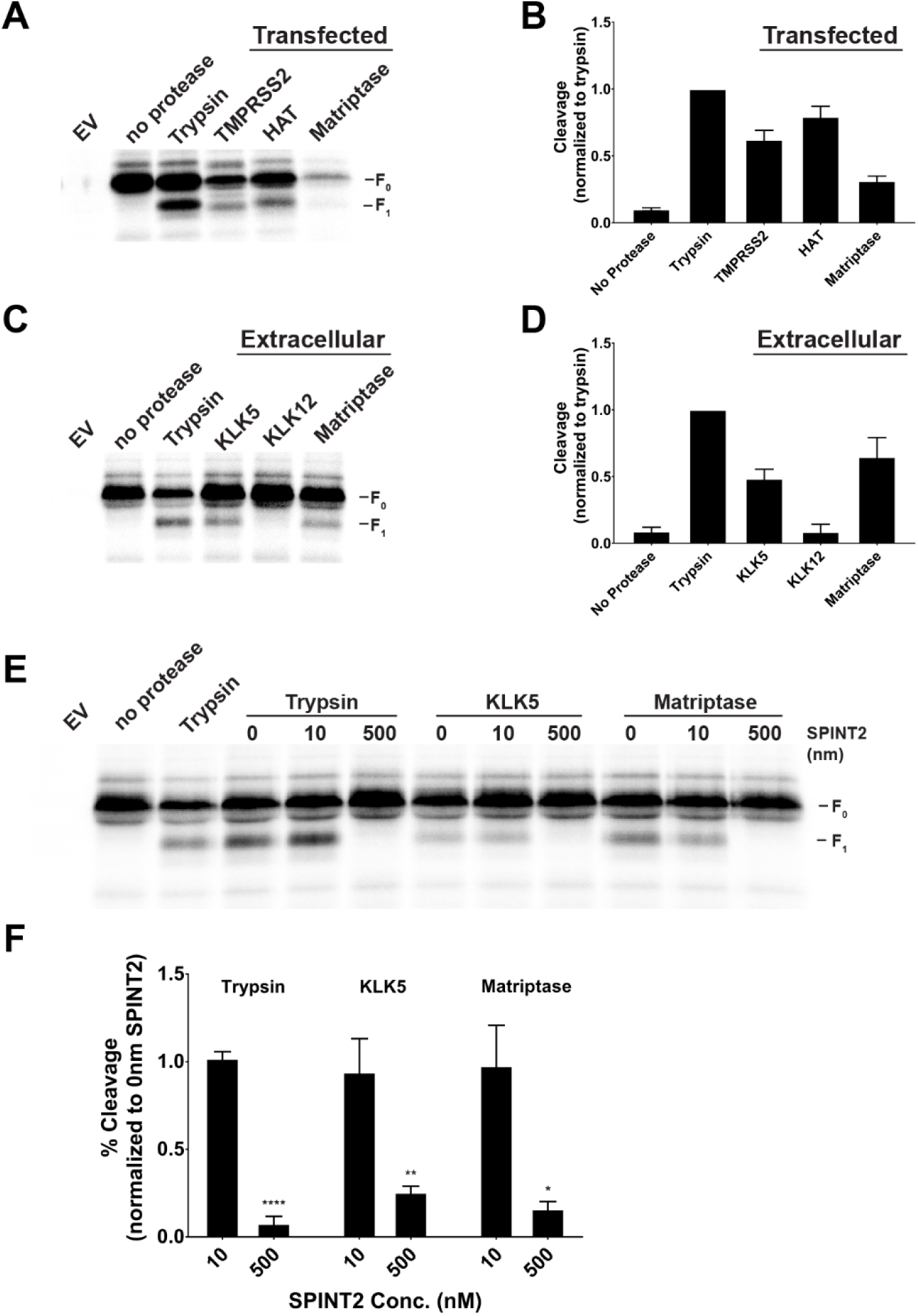
TMPRSS2, HAT, matriptase and KLK5 cleave HMPV F and SPINT2 is able to prevent cleavage by exogenous proteases. HMPV F was either expressed alone or co-transfected with protease and allowed to express for ∼ 18hours. Cells were then metabolically starved of cysteine and methionine followed by radioactive S35 labeling of protein for 4 hours in the presence of TPCK-trypsin or specified protease. SPINT2 treated proteases were incubated at room temperature for 10 minutes and placed onto cells for 4 hours. Radioactive gels were quantified using ImageQuant software with percent cleavage equal to 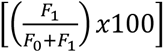. A and B) Co-transfected proteases TMPRSS2, HAT and matriptase are able to cleave HMPV F (n=4) while C and D) exogenous proteases KLK5 and matriptase but not KLK12 are able to cleave HMPV F (n=5). E and F) SPINT2 prevented cleavage of HMPV F by trypsin, KLK5 and matriptase at nm concentrations demonstrated by the loss of the F_1_ cleavage product (n=3). Statistical analysis was performed using a one-way ANOVA followed by a student’s t-test with a bonferroni multiple comparisons test correction. P<0.05 *, P<0.005 **, P<0.005 ***, P<0.001 ****. N values represent independent replicates for each treatment group. Error bars represent SD.

### SPINT2 inhibits HA mediated cell-cell fusion

The above presented biochemical experiments demonstrate that SPINT2 is able to efficiently inhibit proteolytic cleavage of HMPV F and several influenza A HA subtypes by a variety of proteases. For a functional analysis to determine whether SPINT2 inhibition prevents cells to cell fusion and viral growth, we examined influenza A infection in cell culture. While HMPV is an important human pathogen, Influenza grows significantly better in cell culture compared with HMPV. First, we tested whether cleavage inhibition by SPINT2 resulted in the inhibition of cell-cell fusion. As described above, matriptase and KLK5 were pre-incubated with 10nM and 500nM SPINT2 and subsequently added to VERO cells expressing A/CA/04/09 (H1N1) HA or A/Shanghai/2/2013 (H7N9) HA. Cells were then briefly exposed to a low pH buffer to induce fusion and subsequently analyzed using an immune fluorescence assay. When matriptase and KLK5 were tested with 10nM SPINT2 and incubated with VERO cells expressing H1N1 HA, we still observed syncytia formation (Figure 4A). However, 500mM SPINT2 resulted in the abrogation of syncytia formation triggered by cleavage of the respective HA by matriptase and KLK5. We made the same observation when we tested KLK5 and H7N9 HA (Figure 4B). Matriptase-mediated H7N9 HA syncytia formation was inhibited by the addition of 10nM SPINT2 (Figure 4B). To ensure that cell-cell fusion inhibition is a result of HA cleavage inhibition through SPINT2 but not a side effect of SPINT2 treatment *per se* we expressed A/Vietnam/1204/2004 (H5N1) HA in VERO cells and treated them with the inhibitor. H5N1 HA possesses a HPAI cleavage site and is cleaved intracellularly by furin during its maturation process. SPINT2 does not inhibit furin and is not able to cross cell membranes. Thus, SPINT2 can not interfere with the proteolytic processing of H5N1 HA and therefore this control allows to examine whether SPINT2 interferes with cell-cell fusion. Figure 4C shows that H5N1 HA forms large syncytia in the absence of SPINT2 as well as in the presence of 500 nM SPINT2. Hence, we conclude that SPINT2 does not have a direct inhibitory effect on cell-cell fusion.

**Figure 4:**
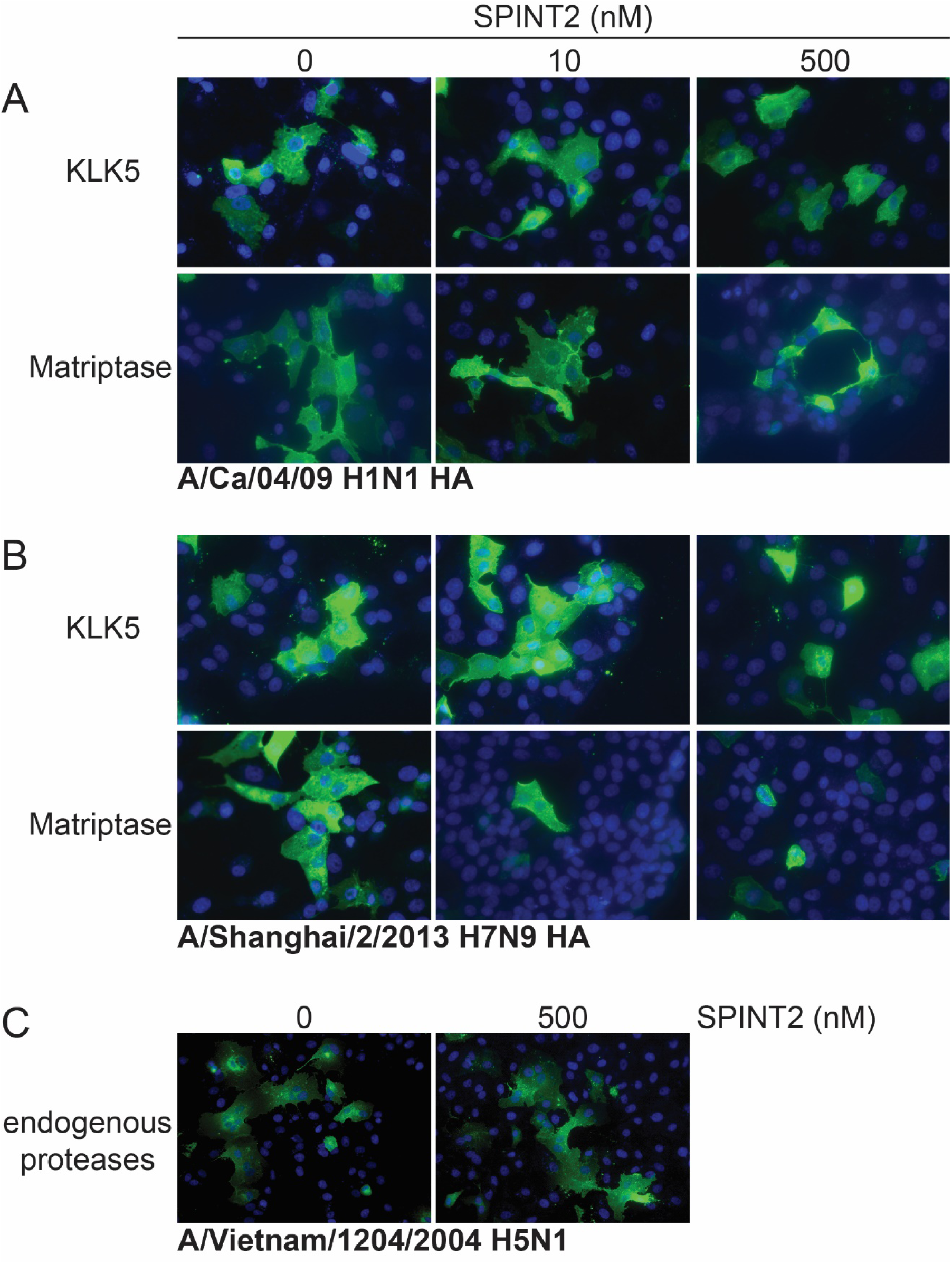
SPINT2 inhibits HA-mediated cell-cell fusion. VERO cells were transfected with plasmids encoding for (A) A/CA/04/09 H1N1 HA or (B) A/Shanghai/2/2013 H7N9 HA and allowed to express the protein for ∼18 hours. Recombinant matriptase and KLK5 were incubated with different SPINT2 concentrations for 15 minutes and then added to the HA-expressing cells for 3 hours. After 3 hours the cells were briefly treated with cell fusion buffer at pH5, washed, supplemented with growth medium and returned to the incubator for 1 hour to allow fusion. HA protein was detected using HA-specific primary antibodies and a secondary fluorogenic Alexa488 antibody. Nuclei were stained using DAPI. Magnification 40x. (C) VERO cells expressed A/Vietnam/1204/2004 H5N1 HA that was cleaved during its maturation process in the cell. SPINT2 was added at 0 nM or 500 nM at the time of transfection. Magnification 25x.

### SPINT2 reduces viral growth in cell culture

To understand whether SPINT2 was able to inhibit or reduce the growth of virus in a cell culture model over the course of 48 hours we transfected cells with human TMPRSS2 and human matriptase, two major proteases that have been shown to be responsible for the activation of distinct influenza A subtype viruses. TMPRSS2 is essential for H1N1 virus propagation in mice and plays a major role in the activation of H7N9 and H9N2 viruses (42–44). Matriptase cleaves H1N1 HA in a sub-type specific manner, is involved in the *in vivo* cleavage of H9N2 HA and our results described above suggest a role for matriptase in the activation of H7N9 (19, 44). At 18 hours post transfection we infected MDCK cells with A/CA/04/09 (H1N1) at a MOI of 0.1 and subsequently added SPINT2 protein at different concentrations. Non-transfected cells served as a control and exogenous trypsin was added to facilitate viral propagation. The supernatants were harvested 48 hours post infection and viral titers were subsequently analyzed using an immuno-plaque assay. SPINT2 initially mitigated trypsin-mediated growth of H1N1 at a concentration of 50nM and the extent of inhibition slightly increased with higher concentrations (Figure 5A, Table 3A). The highest tested SPINT2 concentration of 500nM reduced viral growth by about 1 log (Figure 5A, Table 3A). We observed a similar pattern with cells transfected with human matriptase (Figure 5B, Table 3A). Growth inhibition started at a SPINT2 concentration of 50nM and with the application of 500nM growth was reduced by approximately 1.5 logs (Figure 5B, Table 3A). When we infected cells expressing TMPRSS2 with H1N1 and added SPINT2, viral growth was significantly reduced at a concentration of 150nM. Addition of 500nM SPINT2 led to a reduction of viral growth of about 1.5 logs (Figure 5C, Table 3A). We also tested whether SPINT2 could reduce the growth of a H3N2 virus because it is major circulating seasonal influenza subtype. However, TMPRSS2 and matriptase do not seem to activate H3N2 viruses (19, 45). Hence, trypsin and SPINT2 were added to the growth medium of cells infected with A/X31 H3N2. Compared to control cells without added inhibitor SPINT2 significantly inhibited trypsin-mediated H3N2 growth at a concentration of 50nM (Figure 5D, Table 3A). At the highest SPINT2 concentration of 500nM viral growth was reduced by about 1 log (Figure 5D).

**Figure 5:**
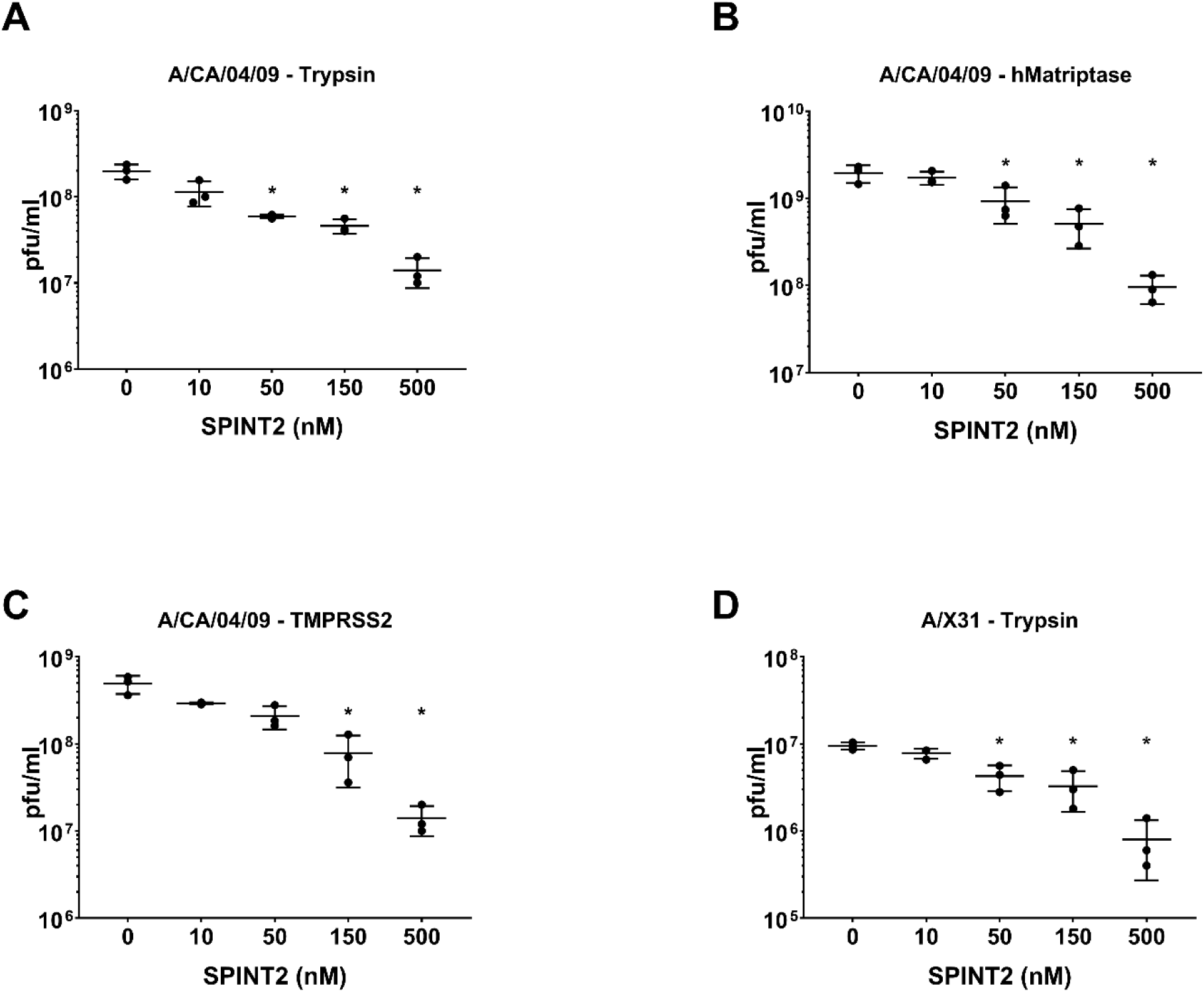
SPINT2 reduces IAV growth in cell culture. MDCK cells were transfected with plasmids encoding for human matriptase or human TMPRSS2 and allowed to express the proteins for ∼18 hours. Cells expressing human matriptase (B) or human TMPRSS2 (C) were then infected with A/CA/04/09 H1N1 at a MOI of 0.1 and different SPINT2 concentration were added to each well. Non-transfected cells to which trypsin was added served as a control (A). (D) MDCK cells were infected with A/X31 H3N2 at an MOI of 0.1 and trypsin was added to assist viral propagation. Different SPINT2 concentration were added as indicated. After 48 hours of infection the supernatants were collected and used for an immuno-plaque assay to determine the viral loads. Experiment was repeated three times and each dot represents the viral titer of a single experiment. Statistical analysis was performed using a non-paired student’s t-test comparing the control (0 h) against the respective sample. Error bars indicate standard deviation. * indicates p = < 0.05. Extended horizontal line within the error bars represents mean value of the three independent replicates.

**Table 3:** Viral titers measured in the infection studies: Table shows the average viral titers and standard deviation calculated from the 3 independent biological replicates depicted in Figures 6 and 7. (A) Titers and standard deviation from infection studies shown in Figure 5. (B) Titers and standard deviation shown in Figure 7.

| SPINT2 | Trypsin - H1N1 |  | Matriptase - H1N1 |  | TMPRSS2 - H1N1 |  | Trypsin - X31 |  |
| --- | --- | --- | --- | --- | --- | --- | --- | --- |
| (nM) | Average pfu/ml | StDev | Average pfu/ml | StDev | Average pfu/ml | StDev | Average pfu/ml | StDev |
| 0 | 1.99E+08 | 3.91E+07 | 1.95E+09 | 4.39E+08 | 4.90E+08 | 1.16E+08 | 9.47E+06 | 9.02E+05 |
| 10 | 1.14E+08 | 3.70E+07 | 1.73E+09 | 2.95E+08 | 2.93E+08 | 7.57E+06 | 7.80E+06 | 1.04E+06 |
| 50 | 5.93E+07 | 3.06E+06 | 9.24E+08 | 4.16E+08 | 2.08E+08 | 6.16E+07 | 4.27E+06 | 1.40E+06 |
| 150 | 4.60E+07 | 8.72E+06 | 5.09E+08 | 2.44E+08 | 7.80E+07 | 4.65E+07 | 3.27E+06 | 1.62E+06 |
| 500 | 1.40E+07 | 5.29E+06 | 9.53E+07 | 3.43E+07 | 1.40E+07 | 5.29E+06 | 8.00E+05 | 5.29E+05 |

**B**
| SPINT2 | Trypsin - H1N1 |  |
| --- | --- | --- |
| (nM) | Average pfu/ml | StDev |
| 0 | 1.88E+08 | 1.06E+07 |
| 500 | 2.80E+07 | 1.00E+07 |
| 500 24hpi | 4.07E+07 | 2.00E+07 |

We also examined the effect of SPINT2 inhibition of HMPV spread over time. VERO cells were infected with rgHMPV at MOI 1 and subsequently treated with 500nM of SPINT2 and 0.3µg/mL of TPCK-trypsin. Every 24 hours, cells were scraped and the amount of virus present was titered up to 96hpi with SPINT2 and trypsin replenished daily. We find that un-treated cells are infected and demonstrate significant spread through 96hpi. Conversely, cells infected and treated with SPINT2 had no detectible virus up to 48hpi and very minimal virus detected at 72 and 96hpi, demonstrating that SPINT2 significantly inhibition HMPV viral replication and spread (Figure 6).

**Figure 6:**
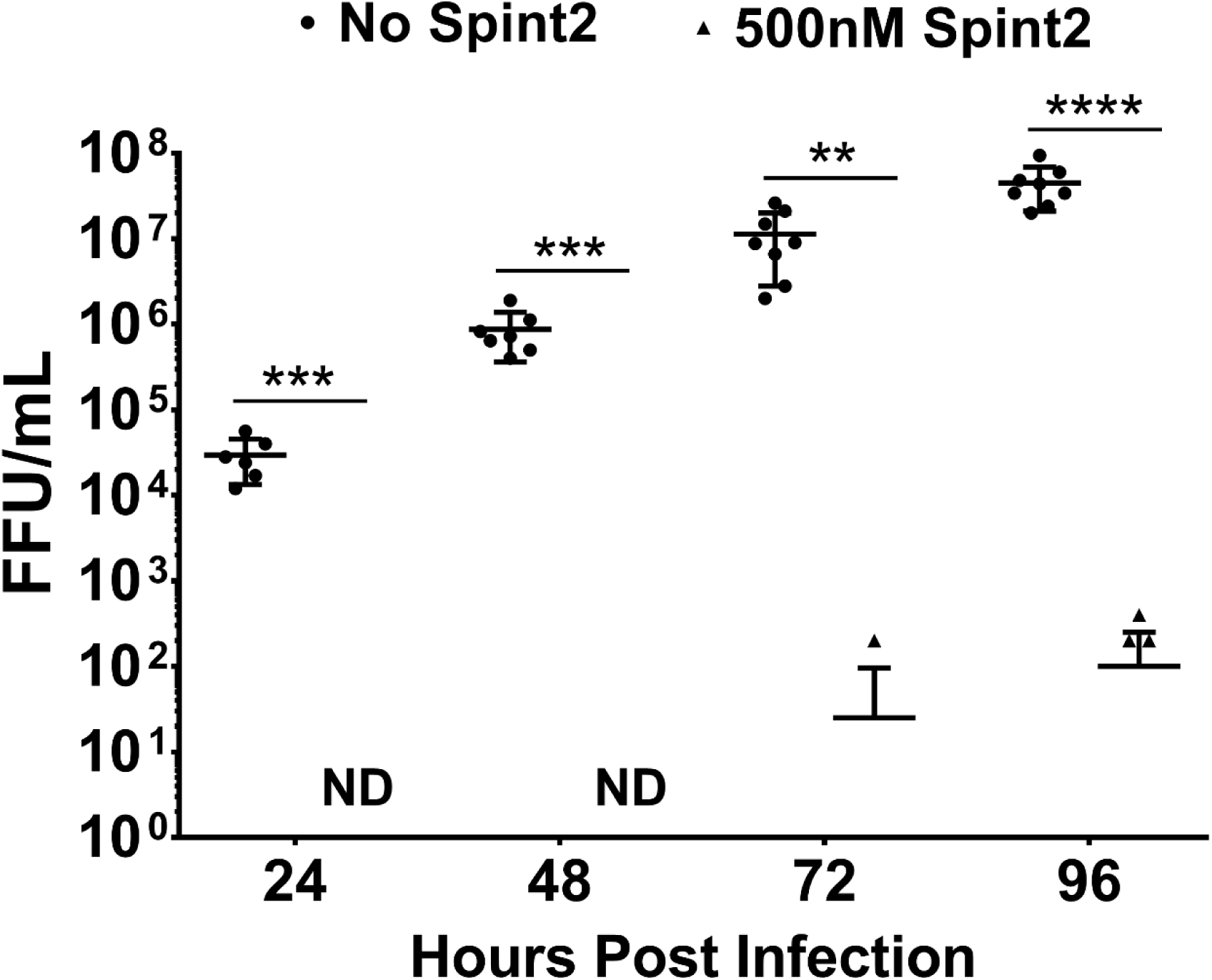
SPINT2 mitigates the spread of HMPV in cell culture. VERO cells were infected with a MOI 1 of rgHMPV. SPINT2 and TPCK-trypsin were added at 500nM and 0.3µg/mL respectively and spread was monitored up to 96hpi. Cells not treated with SPINT2 demonstrated significant spread up through 96hpi whereas treated samples did not show any detectable virus up to 48hpi. However, there was minimal detected virus at 72 and 96hpi. Error bars indicate standard deviation of 4 independent samples completed in duplicate (all data points plotted within the graph). Statistical analysis was performed using a student’s t-test. P <0.05 *, P <0.005 **, P <0.005 ***, P <0.001 ****. ND indicates that the sample was below the limit of detection.

Antiviral therapies are often applied when patients already show signs of disease. Therefore, we tested if SPINT2 was able to reduce viral growth when added to cells 24 hours after the initial infection. Cells were infected with 0.1 MOI of A/CA/04/09 (H1N1) and trypsin was added to promote viral growth. At the time of infection, we also added 500nM SPINT2 to one sample. A second sample received 500nM SPINT2 24 hours post infection. Growth supernatants were harvested 24 hours later, and viral growth was analyzed. We found that viral growth was significantly reduced by regardless whether SPINT2 was added at the time of infection or 24 hours later (Figure 7, Table 3B).

**Figure7:**
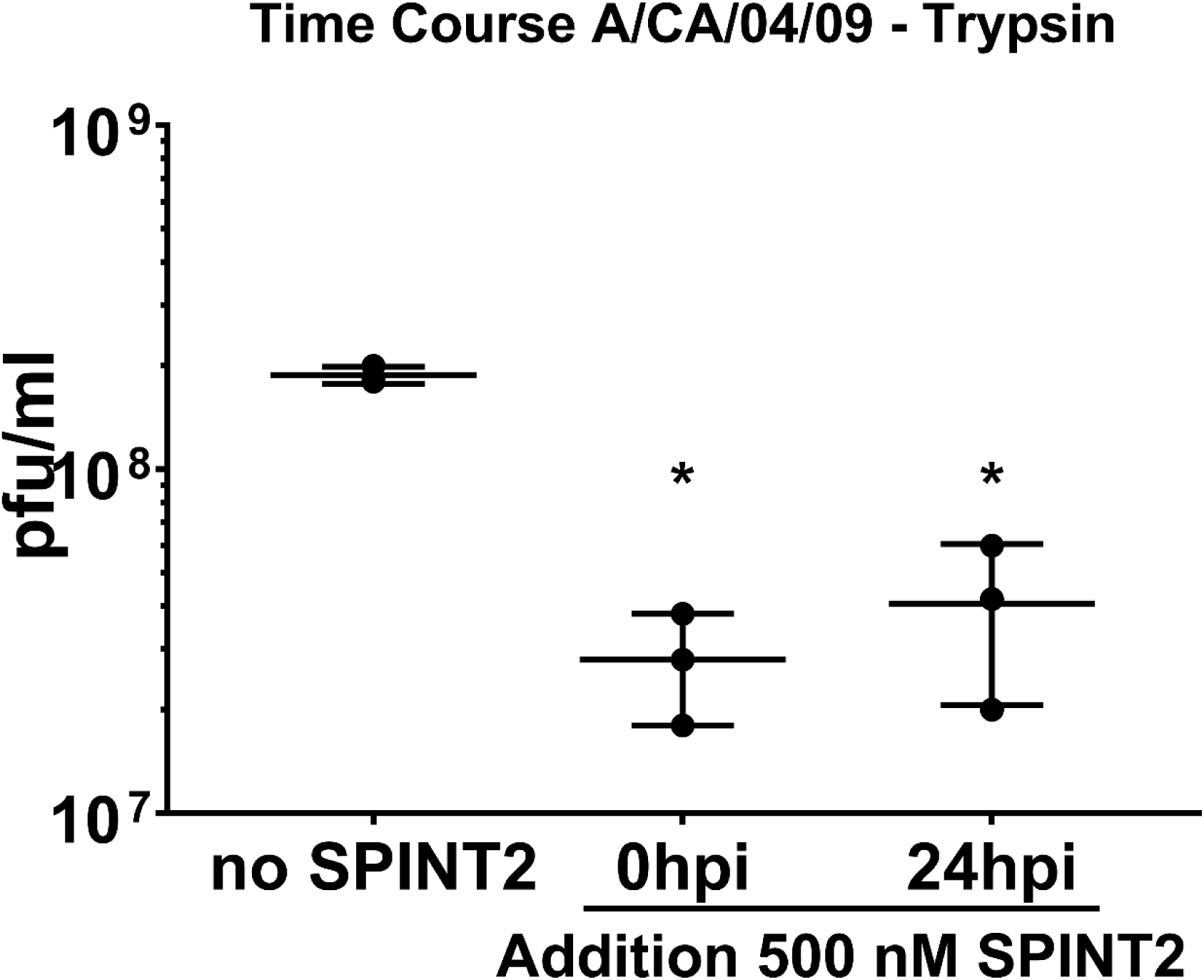
SPINT2 reduces viral growth when added 24 hours post infection. MDCK cells were infected with A/CA/04/09 H1N1 at a MOI of 0.1 and trypsin was added. 500nM SPINT2 were added at the time of infection or 24 hours post infection. Supernatants were collected 48 hours post infection and used for an immuno-plaque assay to determine the viral titers. Statistical analysis was performed using a non-paired student’s t-test comparing the control (0 h) against the respective sample. Error bars indicate standard deviation. * indicates p = < 0.001. Extended horizontal line within the error bars represents mean value of the three independent replicates.

## Discussion

Influenza A virus has caused four pandemics since the early 20^th^ century and infects millions of people each year as seasonal ‘flu, resulting in up to 690,000 deaths annually (5). Vaccination efforts have proven to be challenging due to the antigenic drift of the virus and emerging resistance phenotypes (46). Moreover, the efficacy of vaccines seems to be significantly reduced in certain high-risk groups (47). Prevalent antiviral therapies to treat influenza A virus-infected patients such as adamantanes and neuraminidase inhibitors target viral proteins but there is increasing number of reports about circulating influenza A subtypes that are resistant to these treatments (7). HMPV causes infections in the upper and lower respiratory tract expressing very similar symptoms as influenza infections and resulting in significant morbidity and mortality (11, 12). The most susceptible groups are young children, older adults and people that are immunocompromised (13–15). Currently, there are not treatment against HMPV infections available. In this study we focused on a novel approach that uses antiviral therapies targeting host factors rather than viral proteins offering a more broad and potentially more effective therapeutic approach (24). We demonstrate that SPINT2, a potent inhibitor of serine-type proteases, is able to significantly inhibit cleavage of HMPV F and HA, impair HA-triggered fusion of cells and hence, reduce the growth of various influenza A strains in cell culture.

We assessed cleavage of the HMPV fusion protein in vitro using a peptide cleavage assay modified from previously work on other viral fusion proteins (38, 48). The HMPV F peptide was cleaved by trypsin, plasmin, matriptase and KLK5 but was unable to be cleaved by KLK12. To confirm these findings in a system in which the entire HMPV F protein was subject to cleavage, we co-expressed the fusion protein with TMPRSS2, HAT and matriptase proteases, or treated F with exogenous proteases KLK5, KLK12 and matriptase. These findings are the first to identify proteases besides trypsin and TMPRSS2 that are able to cleave HMPV F. In addition, HMPV appears to utilize many of the serine proteases that influenza uses for HA processing and therefore, offers strong potential for an antiviral target.

SPINT2 demonstrates greater advantage over other inhibitors of host proteases such as e.g. aprotinin that was shown to be an effective antiviral but also seemed to be specific only for a subset of proteases (25). It can be argued that a more specific protease inhibitor which inhibits only one or very few proteases might be more advantageous because it may result in less side effects. With respect to influenza A infections, TMPRSS2 could represent such a specific target as it was shown to a major activating proteases for H1N1 and H7N9 in mice and human airway cells (42, 44, 49–51). However, there is no evidence that that application of a broad-spectrum protease inhibitor results in more severe side effects than a specific one as side effects may not be a consequence of the protease inhibition but the compound itself may act against different targets in the body. The reports demonstrating that TMPRSS2 is crucial for H1N1 and H7N9 virus propagation in mice and cell culture suggest that it also plays a major role in the human respiratory tract. So far, however, it is unclear whether the obtained results translate to humans and other studies have shown that for example human matriptase is able to process H1N1 and H7N9 (19, 52).

Our peptide assay suggests that SPINT2 has a wide variety of host protease specificity. With the exception of plasmin, all the tested proteases in combination with peptides mimicking the cleavage site of different HA subtypes expressed IC_50_ values in the nanomolar range. Interestingly, the IC50 values obtained for cleavage inhibition of HMPV F were substantially lower, in the picomolar range. This suggests that the HMPV cleavage may be more selectively inhibited by SPINT2. However, the western blot data showed that addition of the lowest concentration (10 nM) of SPINT2 did not result in cleavage inhibition of HMPV F by the tested proteases. Differences in sensitivity of SPINT2 between influenza HA and HMPV require further investigation.

SPINT2 poses several potential advantages over other inhibitors that target host proteases. Cell culture studies showed that, for example, matriptase-mediated H7N9 HA cleavage was efficiently inhibited at a concentration of 10nM SPINT2. In contrast, the substrate range for aprotinin, a serine protease inhibitor shown to reduce influenza A infections by targeting host proteases, seemed to be more limited (25). Other synthetic and peptide-like molecules designed to inhibit very specific serine proteases such as TMPRSS2, TMPRSS4 and TMPRSS11D (HAT) were only tested with those proteases and their potential to inhibit other proteases relevant for influenza A activation remains unclear (53–55). Currently, the most promising antiviral protein inhibitor is camostat which is already approved in Japan for the treatment of chronic pancreatitis (56). Recently, it was demonstrated that camostat inhibited influenza replication in cell culture and prevented the viral spread and pathogenesis of SARS-CoV in mice by inhibition of serine proteases (55, 57). However, camostat was applied prior to the virus infection and it was administered into the mice via oral gavage (57). A previous study showed that SPINT2 significantly attenuated influenza A infections in mice using a concentration that was 40x lower than the described camostat concentration and intranasal administration was sufficient (26). Our current study suggests that SPINT2 is able to significantly inhibit viral spread during an ongoing infection and does not need to be applied prior or at the start of an infection. The mouse study also showed that SPINT2 can be applied directly to the respiratory tract while camostat that is currently distributed as a pill and therefore less organ specific. In addition, camostat is synthetic whereas SPINT2 is a naturally occurring molecule which may attenuate potential adverse effects due to non-native compounds activating the immune system. Future research will be conducted to test if SPINT2 can be applied more efficiently via an inhaler and to explore potential side effects in mice studies. However, when we tested the potential of SPINT2 to inhibit viral replication in a cell culture model we were only able to achieve growth reductions of approximately 1-1.5 logs after 48 hours with a concentration of 500nM SPINT2. One potential explanation is that 500nM SPINT2 was unable to saturate the proteases present in the individual experiments and was not sufficient to prevent viral growth. In addition, the continuous overexpression of matriptase and TMPRSS2 may have produced an artificially high quantity of protein that exceeded the inhibitory capacities of SPINT2. This problem could be solved either by using higher concentration of SPINT2 or by optimizing its inhibitory properties. However, the data also demonstrates that SPINT2 has the ability to inhibit proteases that expressed on the cell surface and that inhibition is not limited to proteases that were added exogenously and pre-incubated with the inhibitor (Figure 5). SPINT2 did not express any cytotoxic effects up to a concentration of 10mM, significantly above the therapeutic dosage required for inhibition. In comparison with other studies, the SPINT2 concentration we used here were in the nanomolar range while other published inhibitors require micromolar concentrations (53–55). However, we believe that future research will allow to fully utilize the potential of SPINT2 as a broad-spectrum antiviral therapy. Wu et al., recently described that the Kunitz domain I of SPINT2 is responsible for the inhibition of matriptase (28). In future studies we will explore whether the inhibitory capabilities of SPINT2 can be condensed into small peptides that may improve its efficacy. Its ability to inhibit a broad range of serine protease that are involved in the activation of influenza A suggest that a SPINT2 based antiviral therapy could be efficient against other pathogens too. TMPRSS2, for example, not only plays a major role in the pathogenesis of H1N1 but is also required for the activation of SARS-CoV and MERS-CoV and HMPV (58, 59). Currently, treatment options for these viruses are very limited and therefore SPINT2 could become a viable option if its potential as an antiviral therapeutic can be fully exploited.

However, while SPINT2 has a therapeutic potential to treat ILIs caused by viruses that require activation by trypsin-like serine proteases it may have its limitation to provide a treatment option for infections caused by influenza HPAI viruses, such as H5N1 (60). These viruses are believed to be activated by furin and pro-protein convertases that belong to the class of subtilisin-like proteases (61). Preliminary data from our lab demonstrated that SPINT2 did not inhibit furin-mediated cleavage of HPAI cleavage site peptide mimics as well as peptides carrying described furin cleavage sites (data not shown). In addition, furin acts intracellularly and we have no evidence that SPINT2 is able to penetrate the cell membrane and thus inhibiting proteases located in intracellular compartments. Therefore, it seems unlikely that SPINT2 is able to inhibit furin in cell culture-based studies or *in vivo* experiments.

In conclusion we believe SPINT2 has potential to be developed into a novel antiviral therapy. In contrast to most similar drugs that are synthetic, SPINT2 is an endogenously expressed protein product that confers resistance to a variety of pathogenic viruses and which can potentially be delivered directly into the respiratory tract as an aerosol. Most importantly, SPINT2 demonstrated the ability to significantly attenuate an ongoing viral infection in cell culture and further research will be conducted to explore the time period during which SPINT2 demonstrates the highest efficacy.

## Materials and Methods

### Cells, plasmids, viruses, and proteins

293T, VERO and MDCK cells (American Type Culture Collection) used for influenza experiments were maintained in Dulbecco’s modified Eagle medium (DMEM) supplemented with 25 mM HEPES (Cellgro) and 10% fetal bovine serum (VWR). VERO cells used for HMPV experiments were maintained in DMEM (HyClone) supplemented with 10% FBS (Sigma). The plasmid encoding A/CA/04/09 (H1N1) HA was generated as described (19). The plasmid encoding for HMPV F S175H434 was generated as described(62). The plasmids encoding for A/Shanghai/2/2013 (H7N9) HA, human TMPRSS2 and human matriptase were purchased from Sino Biological Inc. The plasmid encoding for A/Aichi/2/68 (H3N2) HA was generously donated by David Steinhauer. A/CA/04/09 and A/X31 viruses were propagated in eggs. All recombinant proteases were purchased as described (39).

### Expression and purification of SPINT2

SPINT2 was expressed and purified as described with minor modification (26). In brief, *E. coli* RIL (DE3) arctic express cells (Agilent) were transformed with SPINT2-pSUMO. Cells were then grown in 0.5 L Luria Broth containing 50 µg/ml kanamycin at 37°C. At OD 0.5-0.6 cells were chilled on ice and protein expression was induced with 0.2 mM IPTG. Cells were then grown over night at 16°C. Cells were harvested and protein was purified as previously described (26). SPINT2 protein was eluted by a 1-hour incubation with ULP1-6xHis. Glycerol was added to the eluted SPINT2 protein to a final concentration of 20% and protein aliquots were stored at −80°C. Protein concentration was determined by analyzing different dilutions of SPINT2 on an SDS-PAGE gel along with 5 defined concentrations of BSA between 100 ng and 1 µg. The gel was then stained with Coomassie, scanned with ChemiDoc Imaging system (Bio-Rad) and bands were quantified using Image Lab software (Bio-Rad). Concentrations of the SPINT2 dilutions were determined based on the BSA concentrations and the final SPINT2 concentration was calculated based on the average of the SPINT2 different dilutions.

### Peptide Assays

Peptide assays were carried out as described (39). The sequence of the HMPV F peptide mimicking the HMPV F cleavage site used in this assay is ENPRQSRFVL including the same N- and C-terminal modifications as described for the HA peptides (39). The V_max_ was calculated by graphing each replicate in Microsoft Excel and determining the slope of the reaction for every concentration (0 nM, 1 nM, 5 nM, 10 nM, 25 nM, 50 nM, 75 nM, 150 nM, 300 nM and 500 nM). The V_max_ values were then plotted in the GraphPad Prism 7 software against the log10 of the SPINT2 concentration to produce a negative sigmoidal graph from which the IC_50_, or the concentration of SPINT2 at which the V_max_ is inhibited by half, could be extrapolated for each peptide protease mixture. Since the x-axis was the log10 of the SPINT2 concentration, the inverse log was then taken for each number to calculate the IC_50_ in nM.

### Cytotoxicity assay

The cytotoxicity assay was performed with a cell counting kit-8 (Dojindo Molecular Technologies) according to the manufacturer’s instructions. In brief, approx. 2 × 10^3^ 293T cells were seeded per well of a 96-well plate and grown over night. SPINT2 was added at the indicated concentrations. DMEM and 500 µM H_2_O_2_ were used as a control. 24 hours later 10 µl CKK-8 solution were added to each well and incubated for 1 hour. Absorbance at 450 nm was measured using a SPARK microplate reader (Tecan). Per sample and treatment three technical replicates were used and the average was counted as one biological replicate. Experiment was conducted three times.

### Metabolic protein labeling and immunoprecipitation of HMPV fusion protein

VERO cells were transfected with 2ug of pDNA using Lipofectamine and plus reagent (Invitrogen) in opti-mem (Gibco) according to the manufacturers protocol. The following day, cells were washed with PBS and starved in cysteine/methionine deficient media for 45 min and radiolabeled with 50uC/mL S35-cysteine/methionine for 4 hours. Cells were lysed in RIPA lysis buffer and processed as described previously (22) and the fusion protein of HMPV was immunoprecipitated using anti-HMPV F 54G10 monoclonal antibody (John Williams, U. Pitt). Samples were run on a 15% SDS-PAGE and visualized using the typhoon imaging system. Band densitometry was conducted using ImageQuant software (GE).

### SPINT2 cleavage inhibition by Western blot and cell-cell fusion assay

Experiments were performed as previously described with minor modifications (26). Treatment with trypsin, recombinant matriptase and KLK5 were conducted as described (19, 63). For western blot analysis 293T cells were transfected with Turbofect (Invitrogen) according to manufacturer’s instructions. Cell-cell fusion assays were conducted with VERO cells that were transfected with Lipofectamine according to the manufacturer’s instructions. Antibodies to detect A/CA/04/09 (H1N1) HA (NR28666), A/Aichi/2/68 (H3N2) HA (NR3118),A/Shanghai/2/2013 (H7N9) HA (NR48765) and A/Vietnam/1204/2004 (H5N1) HA (NR2705) were obtained from the Biodefense and Emerging Infections Research Resources Repository. Respective secondary antibodies had an Alexa488 tag (Invitrogen). Western blot membranes were scanned using a ChemiDoc imaging system (Bio-Rad). For quantification the pixel intensity of the individual HA_1_ bands was measured using ImageJ software and cleavage efficiencies were calculated by the following equation: HA_1_ 10 nM or 500 nM SPINT2/HA_1_ 0nM SPINT2 × 100%. Cell-cell fusion assays were carried out as described (19).

### Inhibition of influenza viral infection in cell culture

MDCK cells were seeded to a confluency of about 70% in 6-well plates. One plate each was then transformed with a plasmid allowing for the expression of human matriptase or human TMPRSS2. One plate was transformed with empty vector. 18 hours post transfection cells were infected with the respective egg-grown virus at a MOI of approx. 0.1. Different SPINT2 concentrations were added as indicated. 0.8 nM trypsin was added to the cells transformed with the empty vector. 48 hours post infection supernatants were collected, centrifuged and stored at −80°C. Viral titers were determined using an immune-plaque assay as described (64).

### Inhibition of HMPV viral infection in cell culture

VERO (200,000) cells were plated into 24-well plates. The following day, cells were infected with MOI 1 rgHMPV for 3 hours. Cells were washed with PBS and 500µL OPTI-mem with or without 500nM SPINT2 and 0.3µg/mL of TPCK-trypsin was added and incubated for up to 96 hours. The SPINT2 and trypsin was replenished in new OPTI-mem every 24 hours. For each time point, media was aspirated and 100µL of OPTI-mem was added to cells followed by scraping and flash freezing. These samples were then titered on confluent VERO cells up to 10^-6^ dilution to calculate viral titer. Graph shows 4 independent replicates with internal duplicates plotted as individual points. Some data points not shown are due to sample loss, with a minimum of 6 points per group or 3 independent replicates.

## Acknowledgements

J.T.K. and R.E.D. thank Dr. John Williams from University of Pittsburgh Children’s Hospital for kindly providing the 54G10 anti-HMPV F antibody. This work was funded by National Institutes of Health grant 5R21AI117300.

---

**Supplementary Figure 1:**
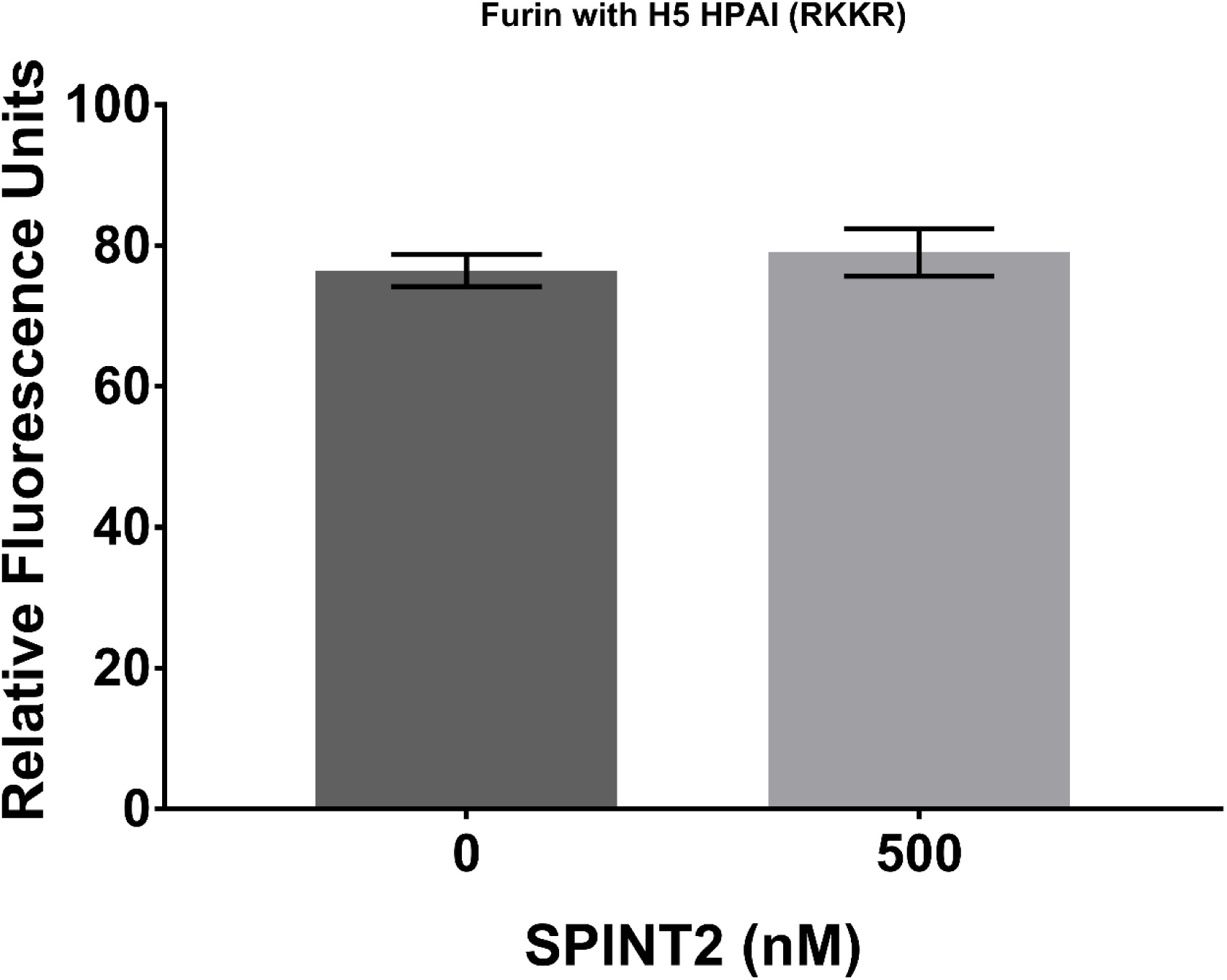
Recombinant furin was incubated with a fluorogenic peptide mimicking the HPAI cleavage site of H5N1 with no SPINT2 present or in the presence of 500 nM SPINT2. Cleavage was monitored by the increase of fluorescence at 390 nm and Vmax values were plotted. n = 3. Error bars represent standard deviation.

**Supplementary Figure 2:**
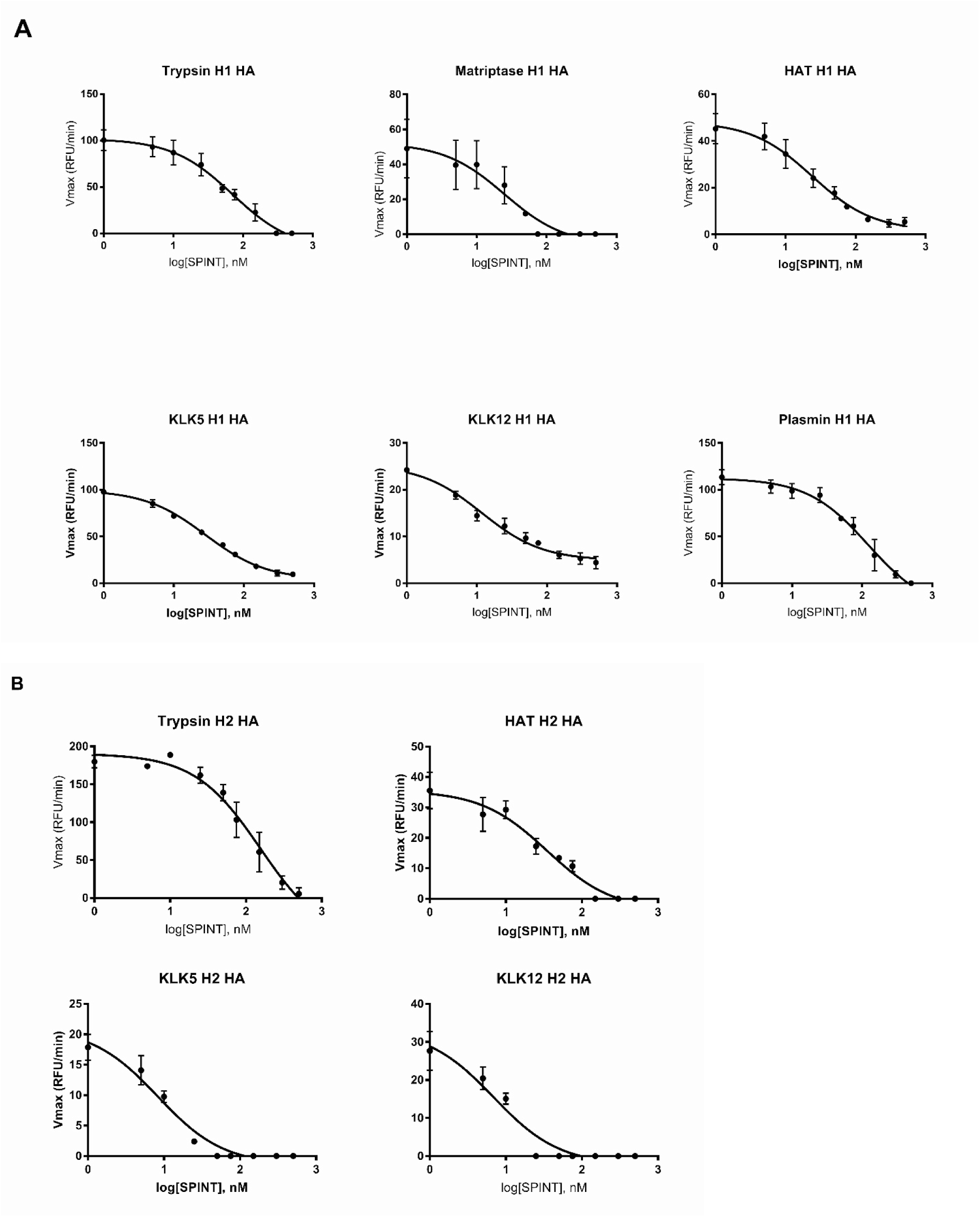
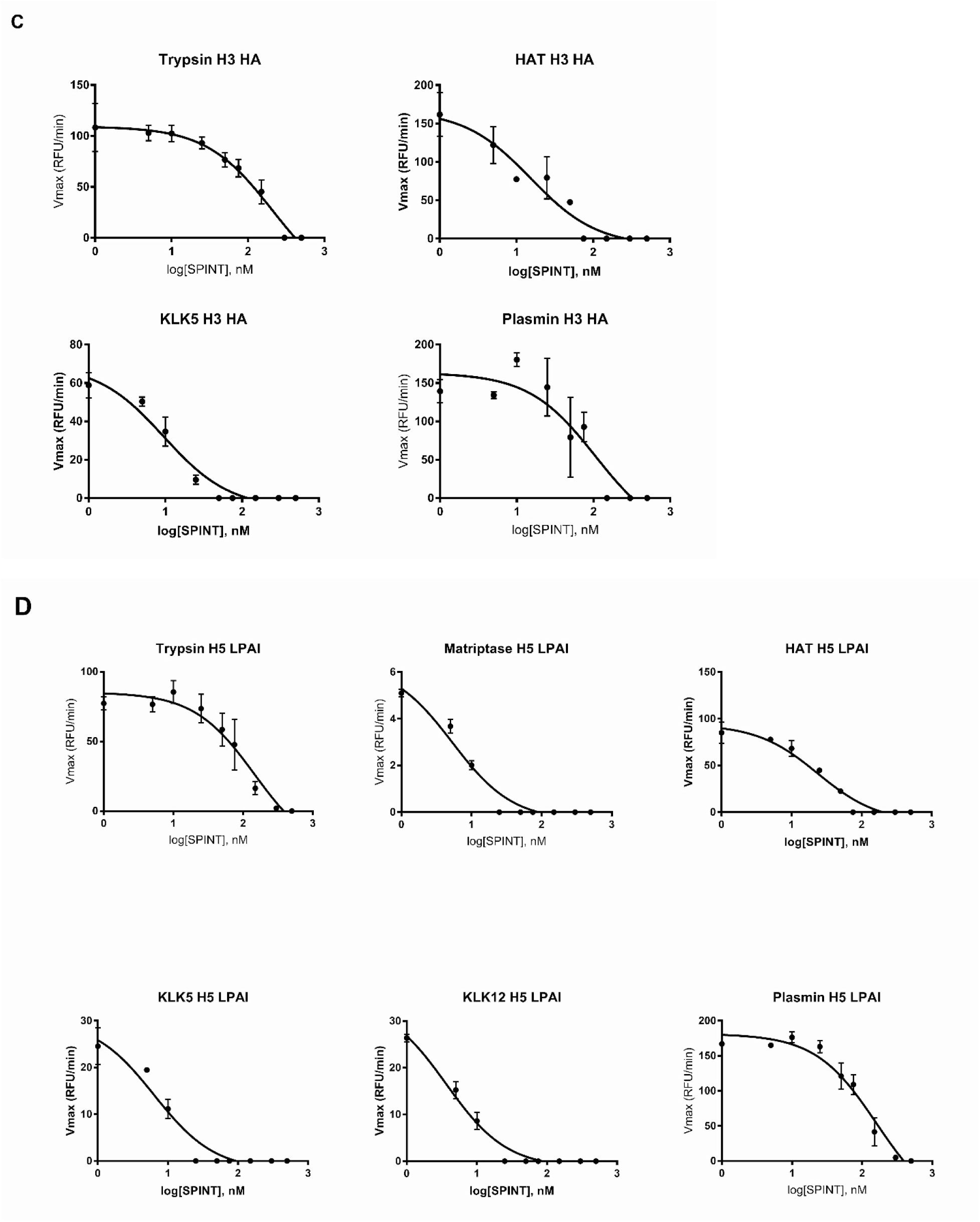
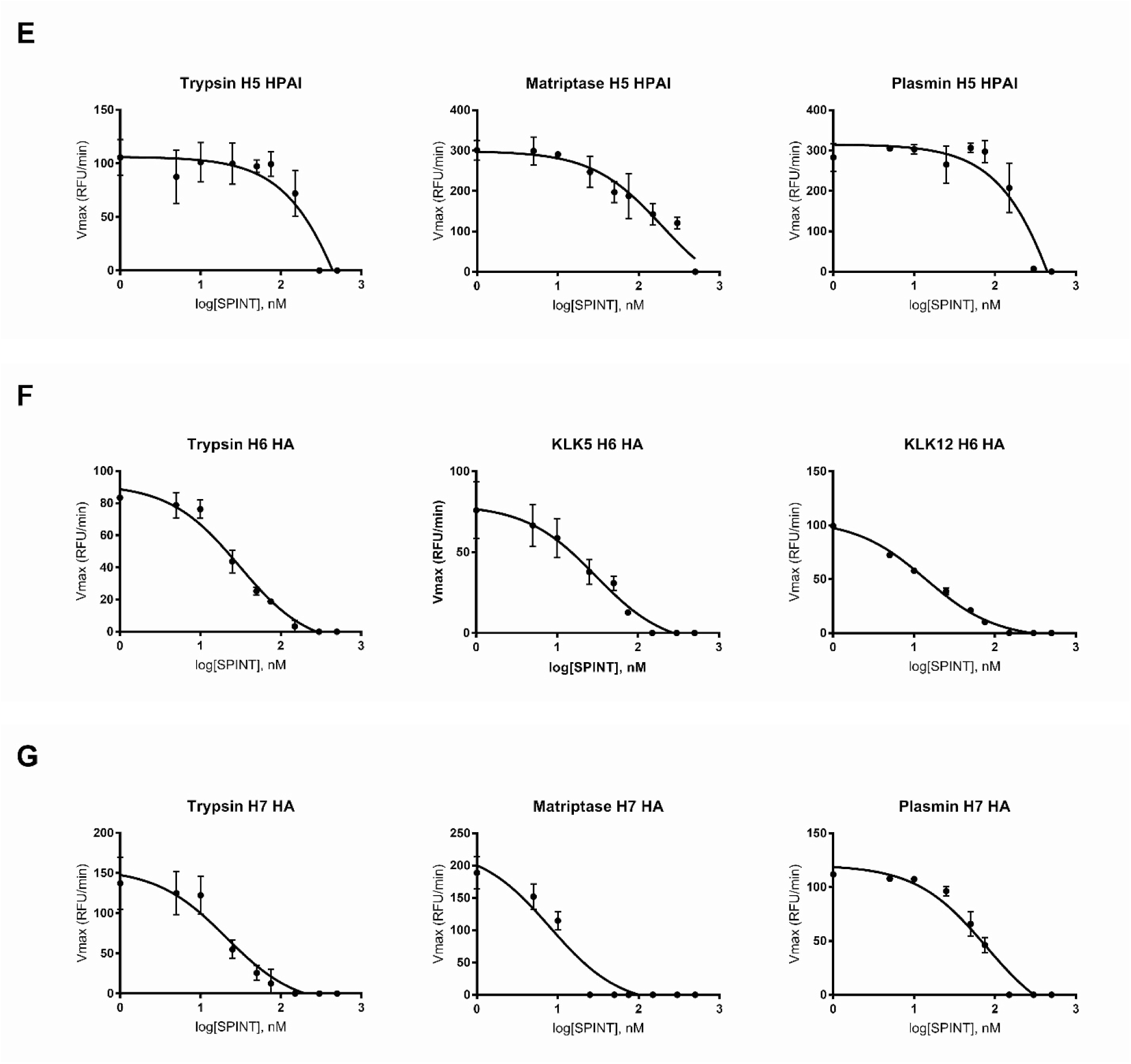
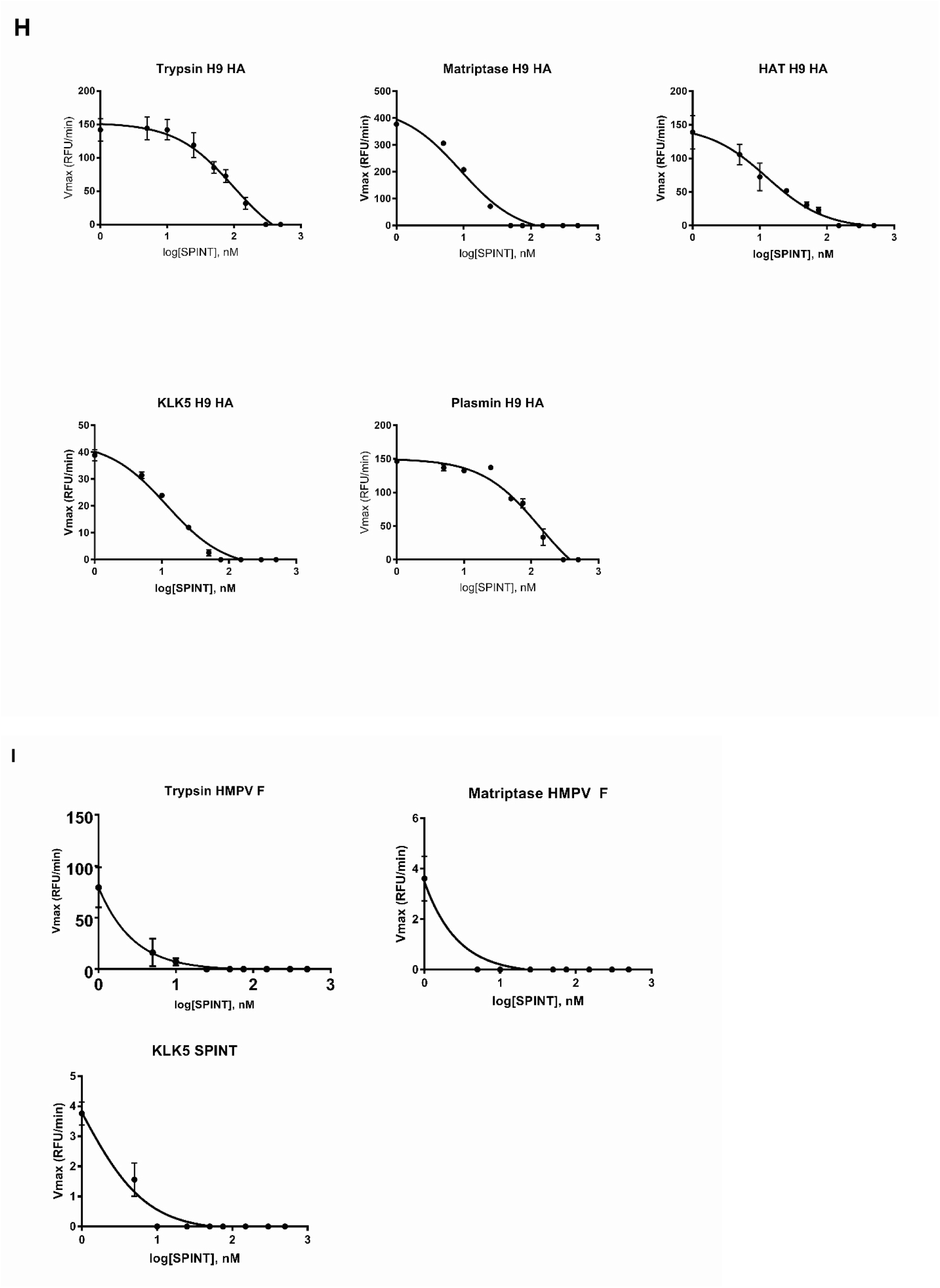
Fluorogenic peptides mimicking the cleavage sites (A) A/CA/04/09 H1N1, (B) A/Japan/305/1957 H2N2 HA, (C) A/Aichi/2/68 H3N2 HA, (D) A/Vietnam/1203/2004 H5N1 LPAI HA, (E) A/Vietnam/1204/2004 H5N1 HPAI HA, (F) A/Taiwan/2/2013 H6N1 HA, (G) A/Shanghai/2/2013 H7N9 HA, (H) A/Hong Kong/2108/2003 H9N2 HA and (I) HMPV F were incubated with the indicated proteases and different SPINT2 concentrations. Cleavage was monitored by the increase of fluorescence at 390 nm and the resulting Vmax values were plotted against the different SPINT2 concentrations on a logarithmic scale (x-axis). Prism 7 software was then used to calculate the IC50 values based on the graphs shown in this figure as described in the “Material and Methods” section. In some cases, the error bars are shorter than the height of the symbol and therefore are not displayed.

